# The Open Perimetry Initiative: a framework for cross-platform development for the new generation of portable perimeters

**DOI:** 10.1101/2021.11.08.467428

**Authors:** Iván Marín-Franch, Andrew Turpin, Paul H Artes, Luke X Chong, Allison M McKendrick, Karam A Alawa, Michael Wall

**Affiliations:** Computational Optometry, Atarfe, Spain; School of Computing and Information Systems, University of Melbourne, Melbourne, Victoria, Australia; Southwest Eye Institute, Tavistock, UK; Eye & Vision Research Group, School of Health Professions, University of Plymouth, Plymouth, UK; School of Medicine (Optometry), Deakin University, Geelong, Australia; Department of Optometry and Vision Sciences, University of Melbourne, Melbourne, Victoria, Australia; Departments of Neurology and Ophthalmology and Visual Sciences, University of Iowa, College of Medicine, Iowa City, Iowa, USA

## Abstract

The Open Perimetry Initiative is fully open source and consists of the Open Perimetry Interface (OPI) and an accompanying package (visualFields) with analytical tools. The OPI package contains an ever-growing number of drivers for commercially available perimeters, head-mounted devices, and virtual reality headsets. The visualFields package contains tools for the analysis and visualization of visual field data, including methods to compute deviation values and probability maps. The use of the OPI and visualFields is shown through a custom static automated perimetry test for the full visual field (up to 50° nasally and 80° temporally) developed with the OPI driver for the Octopus 900 and using visualFields for statistical analysis. Its potential for the development of cross-platform apps for driving and testing portable devices is demonstrated with an OPI driver for an Android-based headset. With more than 55 citations in clinical and translational science as listed in Scopus, this initiative has contributed significantly to expanding the knowledge base in perimetry and clinical vision research at large, and with clinical translation. The continued support of researchers, clinicians, industry, and public institutions are key in transforming perimetry research from closed to open science. The Open Perimetry Initiative provides framework to achieve this.

## Introduction

The advent of new technologies and the development of cross-platform software are paving the way for a new wave of portable devices for visual field testing. The transition from traditional projection perimetry to display-based perimetry requires analysis and adaption of conventional perimetry methods and provides an opportunity to revise and improve.

The Open Perimetry Initiative is an open-source project that started in 2010 with the goal of alleviating the difficulties of using commercial and experimental ophthalmic devices in vision research. The initiative has evolved beyond its original goal to also include features that facilitate the development of new paradigms, standards, and good practices that exploit the technological advantages of portable devices. To maximize accessibility of novel perimetry methods and techniques, implementations using the Open Perimetry libraries are expected to be open source under the GNU General Public License. To avoid the possible legal ramifications, permission from device manufacturers is required before OPI code is used on their commercial instruments.^1^

The key product of the Open Perimetry Initiative is the Open Perimetry Interface (OPI).^1^ The OPI not only provides drivers for an increasing number of ophthalmic devices, but it also sets standards and protocols for the implementation of custom visual field tests so that they can be run seamlessly on different machines, with one implementation for many devices. Furthermore, the OPI can be run in simulation mode, so that new perimetric procedures can be implemented, debugged, and assessed before they are ported to the actual test device. The second key product of the Open Perimetry Initiative is visualFields.^2^ This R package^3^ is a tool for the statistical analysis and visualization of perimetry results. Until now, the OPI and visualFields solutions have been developed largely independently from one another. Recent developments in R, in particular the shiny package^4^ (https://shiny.rstudio.com), have made it possible to easily develop cross-platform applications with graphical interfaces that integrate the OPI and visualFields software.

The purpose of this paper is to describe the recent advances in both the OPI and visualFields and to showcase their use in conventional perimeters as well as new devices such as tablets, phones, and virtual reality headsets. In addition to the development of novel paradigms, methods, and software, the optical properties need to be characterized on these new devices, including resolution and the effects of chromatic and achromatic aberrations. It is likewise necessary to develop adequate methods for calibration and compensation of refractive errors. Considerations of optical characterization, calibration, and limitations are beyond the scope of this manuscript.

## Methods

The OPI has drivers for the Octopus 900 perimeter (Haag-Streit AG, Köniz, Switzerland), the Compass microperimeter (Centervue, Padova, Italy), and the AP-7000 perimeter (Kowa, Torrance, CA, USA). New drivers have now been incorporated for the IMO (CREWT Medical Systems, Tokyo, Japan) and for the Google Cardboard, Google Daydream, and other Android-compatible head-mounted displays. Drivers for devices such as the VIVE Pro Eye (HTC Corporation, Taoyuan City, Taiwan) and AVA Advanced Vision Analyzer (Elisar, India) are being developed.

### OPI conventions and standards

The OPI implementations and drivers follow conventions and standards that not only accelerate software development, but also enable the creation of custom tests, perimetric algorithms, and procedures. These can all be used with different computer operating systems, programming languages, and with different commercial and experimental perimeters. The OPI commands are described more fully elsewhere^1^ but are listed here for completeness:

- opiInitialize(): open connection with the perimeter and ready it,
- opiQueryDevice(): get information about the perimeter,
- opiSetBackground(): set the background color, luminance, fixation point, etc.,
- opiPresent(): Present a stimulus and return observer’s response, and
- opiClose(): close the connection with the perimeters

There are three distinct types of visual field stimuli that can be presented with the opiPresent() command: static, temporal, and kinetic. The static type can be used for conventional static automated perimetry.^5,17,18^ The temporal type can be used to generate stimuli that vary over time and that are supported by the underlying hardware, such as the frequency-doubling illusion^19,20^. The kinetic type can be used to present moving stimuli specified according to the nomenclature introduced by Hans Goldmann.^21^ The opiPresent() command returns the response of the observer (whether the response button was pressed or not after a stimulus presentation and the time between onset of stimulus presentation and response), and the pupil position at the time the button was pressed (if hardware allows).

It is possible to interact with the perimeter via the opiPresent() command and build a stimulus-response logic from which complex test procedures can be constructed. Several common procedures are already built into the OPI package:

- MOCS(): The method of constant stimulus or MOCS,^22^
- fourTwo(): The 4-2 dB staircase,^23,24^
- FT(): The full threshold algorithm,^23,24^ and
- ZEST(): The Bayesian QUEST and ZEST algorithms^25^

The implementation of the aforementioned OPI commands and drivers are necessarily different for each different perimeter; however, the specific implementation can be selected with the command opiChoose(). Thus, once a specific perimetry driver —Octopus900, Compass, IMO, Daydream, etc.— has been selected, a dispatcher is set in place so that the same OPI commands listed earlier can be used without change with all supported hardware.

Using the same approach, it is possible to generate graphical user interfaces that are platform-independent (e.g., the R package shiny^4^), which provides a framework for developing interactive web apps. Shiny was used to develop applications to drive the Octopus 900 perimeter and the Google Daydream headset (shown in Results in this manuscript). The Daydream shiny app allows one to configure the device and perimetry test, obtain the luminance profile (i.e., the correspondence between pixel value and physical luminance, of compatible Android phones), manage datasets, run a threshold test on conventional and custom grids of test locations, and generate graphical reports from the results.

### The visualFields analytical tool

The visualFields package has undergone a major revision moving from version 0.6^2^ to version 1.0 introduced here. Its core functionality is the same but the code has been simplified; some dated ideas and methods have been deprecated, and a number of conventions have been adopted for clarity and simplicity. The most visually striking change has been the adoption of Voronoi tessellation^26–28^ for the graphical representation of visual field data and statistics. In addition, an effort has been made to improve its transparency and the reproducibility of its methods. For instance, the SUNY-IU dataset of healthy subjects that was used in the previous version to generate normative values has been incorporated into the package, vfctrSunyiu24d2, along with a function that generates the normative reference values. Thus, the command nvgenerate(vfctrSunyiu24d2) generates pointwise normative values and the command nvgenerate(vfctrSunyiu24d2, method = “smooth“) generates the normative values used in the visualFields 0.6^2^ using the smoothing techniques as those introduced by Heijl and colleagues for the Statpac 2.^29,30^ Normative datasets, vfctrIowaPC26 and vfctrIowaPeri, and reference values generated with the function nvgenerate for the custom tests used to study the advantages of exploring the full visual field are also included in the package.^8–11^

### Data availability

The release version of OPI can be found at https://CRAN.R-project.org/package=OPI. The release version of visualFields can be found at https://CRAN.R-project.org/package=visualFields. The development version of OPI can be found at https://github.com/turpinandrew/OPI. The development version of visualFields can be found at https://github.com/imarinfr/vf1. Most datasets used in this manuscript can be found within the visualFields package. The dataset of healthy subjects for the full visual field can be found in the visualFields package as well as in https://www.sciencedirect.com/science/article/pii/S2352340918311570. The applications developed based on the OPI and visualFields packages and presented in the manuscript, as well as the datasets are available from the corresponding author on reasonable request.

## Results

Figure 1 illustrates the OPI architecture. Once the hardware is selected, the R OPI client dispatches the commands to connect, disconnect, set background and other settings, and present stimuli to the corresponding OPI server (Octopus 900, Compass, IMO, Cardboard, etc.), which ultimately communicates the commands to the hardware. The server then waits for the hardware to send its state or respond (machine initialized, background lit, pupil position, clicker pressed within a time response window after stimulus onset, etc.) and communicates the response to the OPI client. A web app, as the one shown in the top left of Figure 1 in R with the shiny package, can be developed on top of a program logic to run conventional or custom visual field tests on both regular or irregular grids of test locations with the OPI built-in algorithms (ZEST, full threshold, etc.). It also has the capability to run other procedures, including a suprathreshold test due to Aulhorn^5^ or the binocular Esterman test.^6^ Port communication via internet protocol networks and listeners in the program logic allows for separation of the front-end and the OPI client back-end. This way, the web app used as front-end is easily replaceable by any other front-end developed in Java, Python, MATLAB, or any other suitable programming language. Port communication in R was achieved by creating a background R session with the callr package^7^ that runs in parallel. With this architecture, the program logic embedding the OPI client runs in the background while the web app runs in parallel in the foreground.

**Figure 1:**
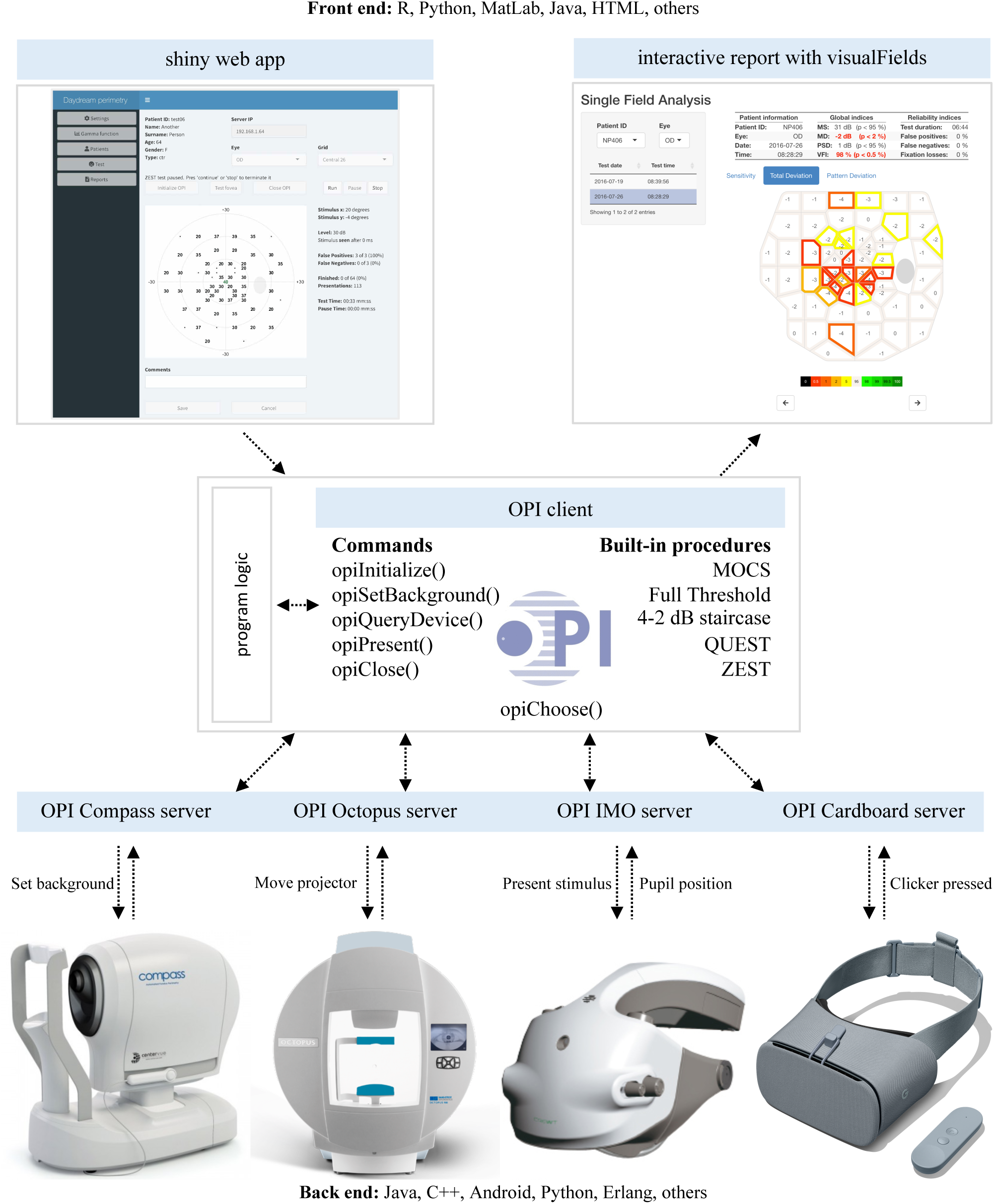
Illustration of the OPI architecture. Top left is a graphical interface generated in shiny for a program logic that can be run on any device at the bottom through the OPI client (center) as it dispatches commands via the OPI servers. Once a suitable dataset of healthy controls has been collected, statistical analyses as the one in the top right can be generated with the visualFields package.

The OPI was used to run a series of tests of the full visual field.^8–11^ A publicly available dataset of 98 eyes of 98 healthy subjects^8^ was used to derive normative values with the visualFields R package.^2^ Figure 2 shows a statistical analysis of the results for a specific full visual field, which consists of the combined analysis of two tests taken on the same day, one for the central visual field and another for the far periphery.^8–11^ The tiles involving each visual field location were obtained using Voronoi tessellation to achieve an efficient representation for both highly irregular grids. Voronoi tessellations are a partitioning of a surface into regions so that the center of each cell is its mean (center of mass). Every point in a given Voronoi polygon is closer to its generating point than to any other cell. We believe this type of graphic best represents the visual area tested, without interpolating between different points The dataset of ocular healthy eyes^8^ is incorporated in the visualFields 1.0 for the central and peripheral tests, vfctrIowaPC26 and vfctrIowaPeri. A script that generates figures for all subjects in those datasets is provided as supplemental material with name vfPlotFullField.r.

**Figure 2:**
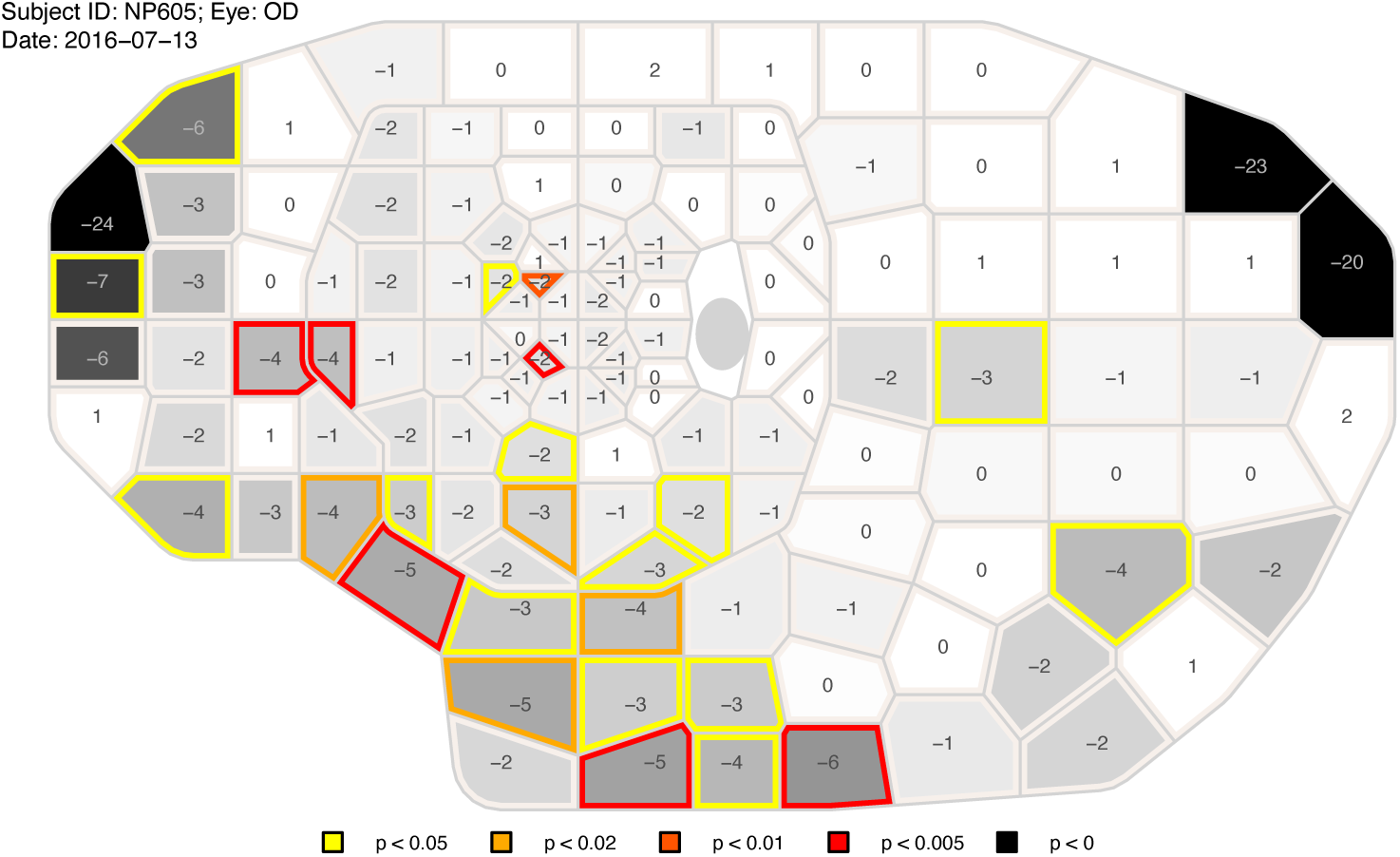
Combined grayscale sensitivity and color-coded total-deviation map. The representation of the full visual field results are composites of two tests taken on the same day: one spanning the central 26° of the visual field and another from 26° to 50° nasally, to 80° temporally, to 50° inferiorly, and to 46° superiorly. The values shown at each location are total deviations, departures in sensitivity from the mean normal sensitivities for age-matched controls. The background grayscale of each tile represents the estimated sensitivity at the corresponding visual field location, where darker means lower sensitivity. Tiles whose border is shown in color are significantly depressed according to the statistical analysis of the total-deviation map. Locations with total-deviation values below the percentile 0.5 from age-corrected mean normal are red, those below the percentile 1 are dark orange, those below the percentile 2 are light orange, and those below the percentile 5 are yellow.

In addition to these single field analyses, the visualFields package also has tools to analyze longitudinal data, including pointwise linear regression, and the permutation of pointwise linear regression, or PoPLR.^12^ Figure 3 shows a brief report (generated with the script vfPoPLRAnalysis.r provided as supplemental material and with the vfpwgSunyiu24d2 dataset^13^). The normative values to obtain total deviation values and probability maps were generated using the dataset, vfctrSunyiu24d2, from a prospective longitudinal study conducted Bloomington, Indiana University (IU) and New York City, State University of New York (SUNY).^2^ The normative values were obtained with the command nvgenerate(vfctrSunyiu24d2).

**Figure 3:**
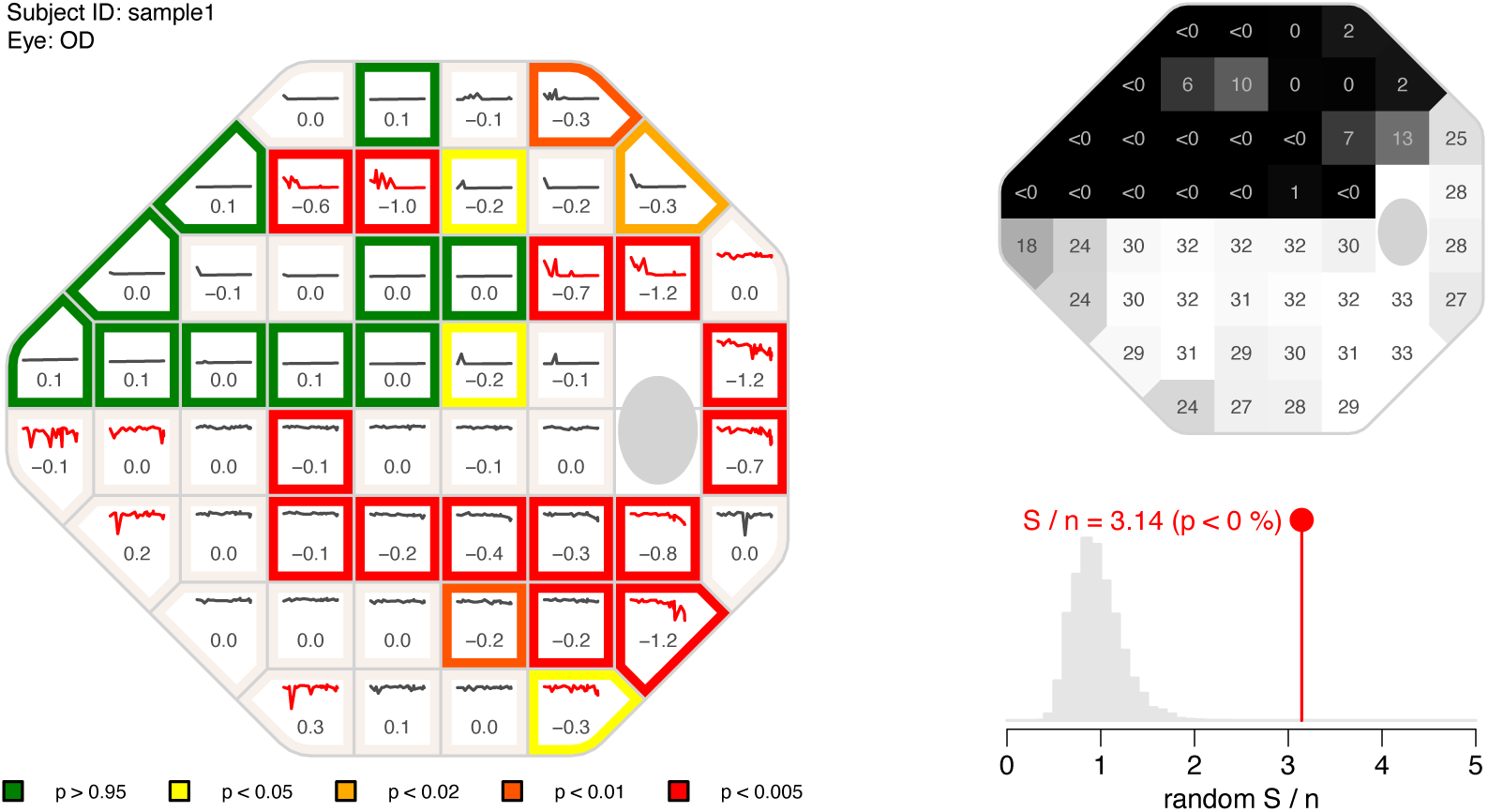
The PoPLR analysis. The left panel shows the slopes at each visual field location obtained with pointwise linear regression of total deviation values over time along with sparklines representing the values over the whole series. The colors at the border of the tiles represent different *p*-values, with red representing a *p*-value lower than 0.5% for the 1-tail *t*-test with the alternative that the slope is negative, dark orange representing *p*-values lower than 1%, light orange representing *p*-values lower than 2%, yellow representing *p*-values lower than 5%, and green representing *p*-values greater than 95%. To identify highly variable series of visual fields, the sparklines are shown in red if the median absolute deviation of the residuals from linear regression were greater than 2 dB. The top right function shows a grayscale map of the baseline sensitivity values (intercept of pointwise regression on sensitivities). The bottom right function shows the histogram of random *S* / *n*, where *n* is the number (52 in this case) of regression analyses performed obtained by permuting the series as part the of PoPLR analysis. The *p*-value for the PoPLR analysis testing whether there is deterioration is shown next to the value of the observed *S* / *n* statistic.

The ZEST algorithm parametrization used to measure the full visual field^8–11^ has been updated and improved for its use with an experimental Daydream OPI perimeter. The specific implementation of the ZEST algorithm has a bimodal prior probability mass function with one peak centered at 0 dB to model sensitivities of damaged locations and another peak that depends on sensitivity estimates in neighboring locations.^14^ Neighboring locations are now defined as locations that share line segments which formed the polygons obtained via Voronoi tessellation. Figure 4 shows the Voronoi tessellation pattern for the 24-2 grid of test locations, with arrows indicating how locations are sampled following a growth algorithm. A conventional 24-2 grid was used instead of the irregular one in Figure 2 for clarity of illustration.

**Figure 4:**
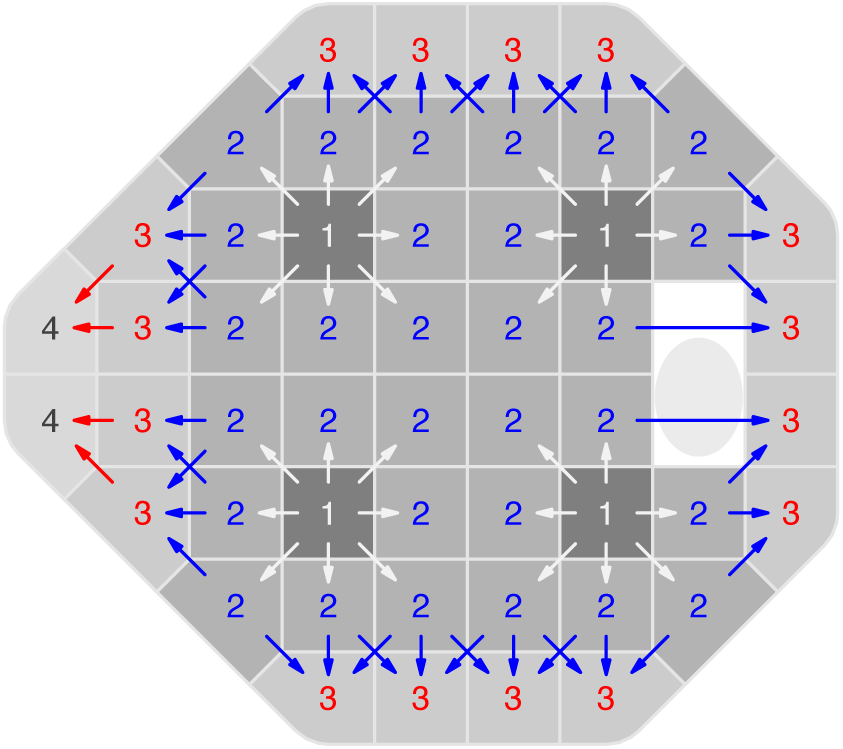
The growth algorithm. The numerals represent the growth pattern wave number of the 52 locations (after excluding the two closest to the blind spot) of the 24-2 regular grid of test locations. Locations corresponding to the primary wave are labeled with the number 1, and locations corresponding to the second, third, and fourth waves are respectively labeled with the numbers 2, 3, and 4. The polygons that determine each location’s wave and neighbors were obtained using Voronoi tessellation and are shown in 4 shades of gray, from dark to light, corresponding to the 4 waves of the growth algorithm.

The growth pattern applied to the 24-2 grid consists of 4 waves and commences at four primary locations (labelled as “1“ in Figure 3). At the beginning, only these four primary locations are tested at random, with the second peak of the bimodal probability mass function at 30 dB. Once a primary location is complete, its neighboring locations that correspond to the second wave (labelled as “2“ in Figure 3) open up for testing, with the second peak centered at the posterior estimated sensitivity threshold of its predecessor. Once a location of the second wave is complete, it propagates its estimated sensitivity threshold to its neighbors that correspond to the third wave (labelled as “2“ in Figure 3) and so on and until all locations have been completed. The growth algorithm should not propagate through the midline since the ganglion cells on the upper and lower hemifields follow different trajectories^15^ and therefore there is no structural correlation between those neighboring visual field locations.

Figure 5 shows a web app developed for the Daydream OPI running the updated ZEST algorithm with an irregular grid of test locations, that corresponds to the central part in the full visual field tests.^8–11^ Although in this application the background was black for the contralateral eye (left half of the screen), the Daydream OPI implementation allows for the presentation of the same background luminance on both eyes.

**Figure 5:**
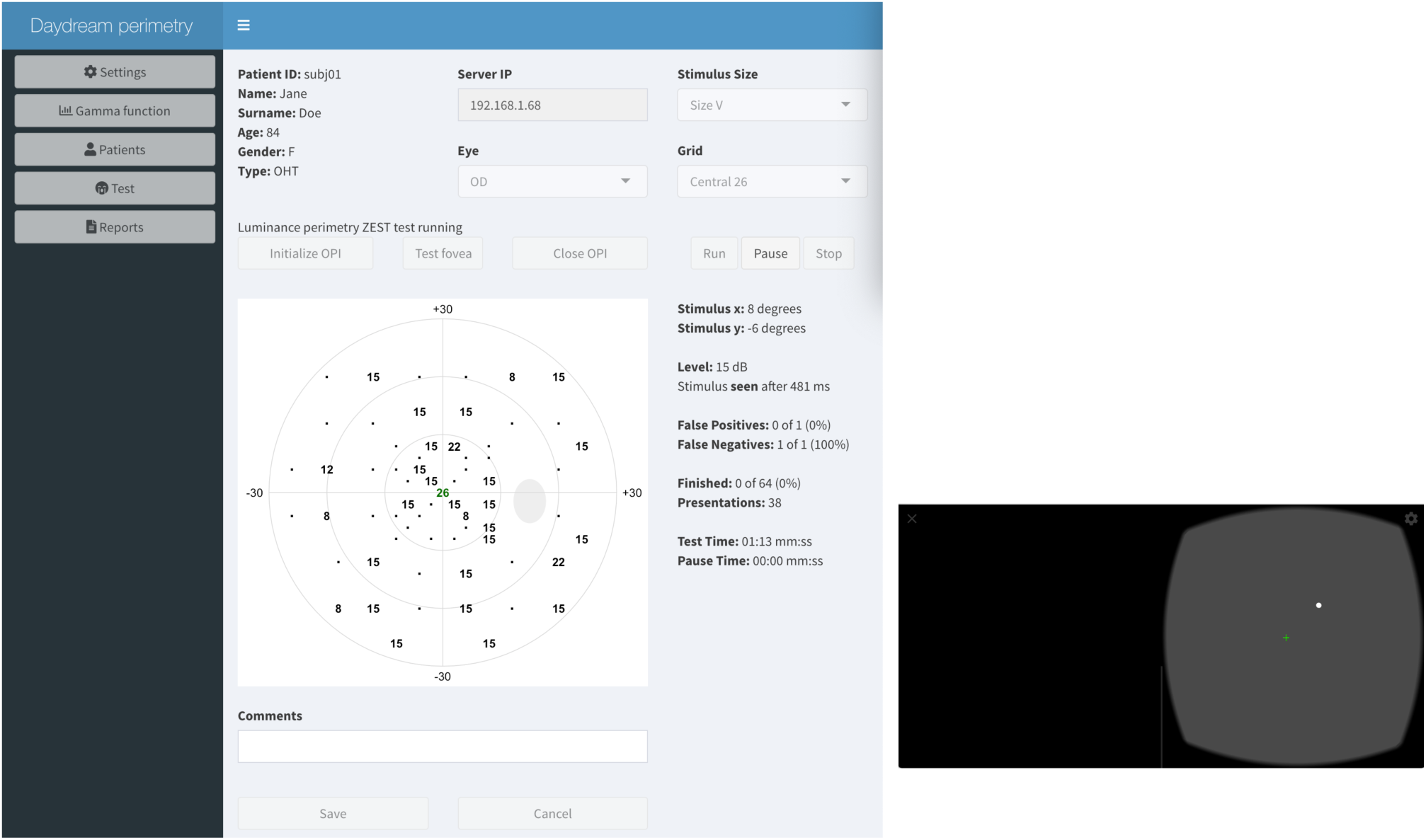
Shiny web app for the OPI Daydream server. The ZEST algorithm for luminance (white-on-white) perimetry for a custom irregular grid of test locations is executed for a (fictitious) patient with ocular hypertension using a web app developed in shiny (left panel). The web app sends commands to an Android Pixel 3 phone (shown in the right panel) to generate white visual stimuli at different intensities and at different distances from the fixation point (green cross). At each presentation, the web app updates and shows the interim results of the test. This screenshot was possible thanks to scrcpy (https://github.com/Genymobile/scrcpy).

## Discussion

The advent of new technologies and the development of cross-platform software are preparing the way for a new wave of portable devices for visual field testing. To lay the groundwork for this new era of perimetry, it is important to build a knowledge base on the strengths and weaknesses of the coming generation of perimeters, as well as revise and accommodate conventional methods and analyses to unveil their full potential. It is essential to do this in a manner that is as transparent and as accessible as possible by adhering to the open science principles (open source, open data, and open access).^16^

The Open Perimetry Initiative and other open-source initiatives that expand perimetry knowledge base and make it freely accessible are essential to equip clinicians and researchers with the necessary tools to objectively assess different products and solutions for the coming generation of portable perimeters. Furthermore, its drivers, applications, and carefully curated datasets collected as a result of these open-source initiatives are beneficial for research groups and institutions, not-for-profit organizations, private entrepreneurs, and eye care at large. The Open Perimetry Initiative continues to have a beneficial impact on perimetry and vision research. As of May 2021, OPI and visualFields related publications received 57 peer-reviewed citations from 12 countries in 4 continents (see Figure 6, data from Scopus). received a total of Its popularity is growing as the yearly average number of citations went from 5.3 before 2018 to 8.3.

**Figure 6:**
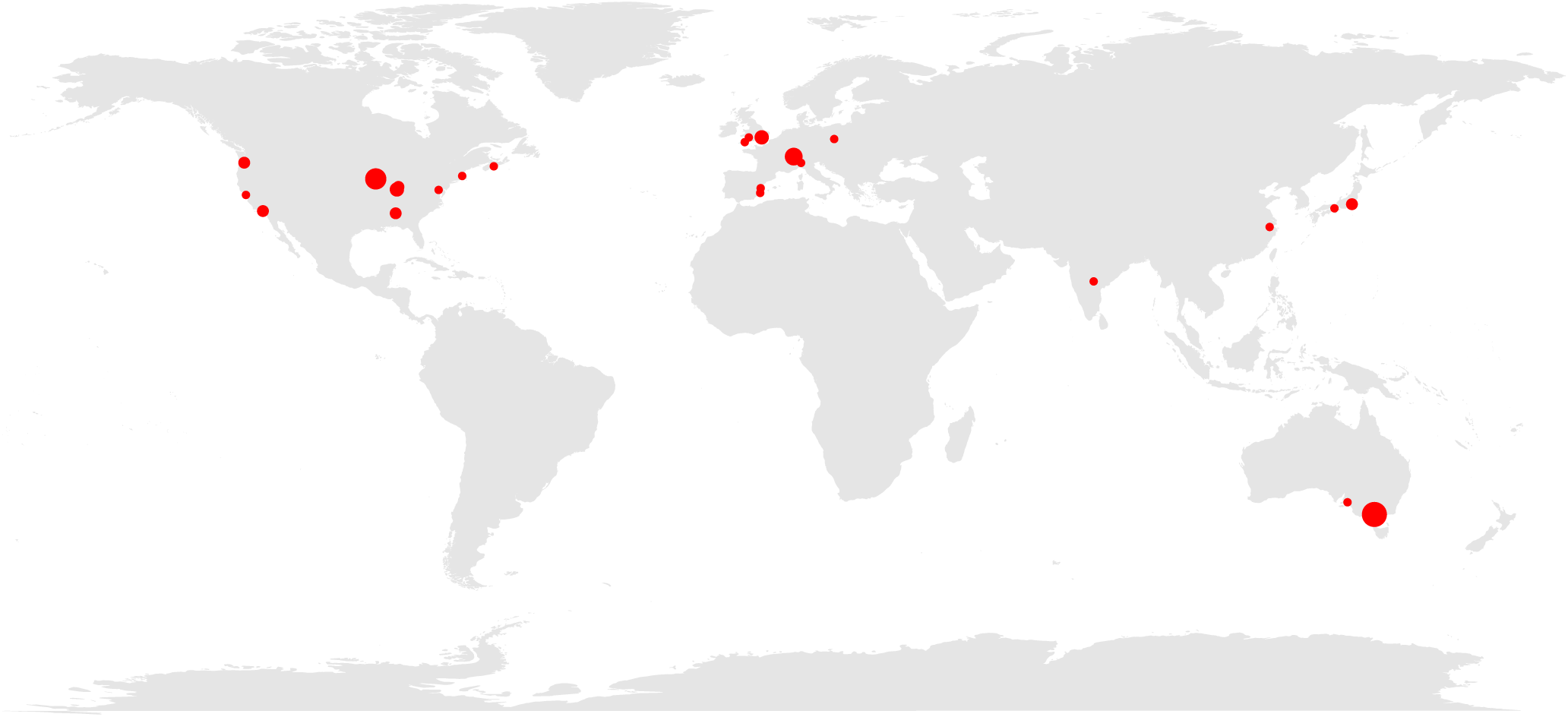
Geographical map of citations to the OPI and the visualFields R package as listed in the ISI Web of Science. The red solid circles demarcate the cities of the first authors’ affiliations. The sizes of the circles represent the number of citations from each city, up to three.

The Open Perimetry Initiative would not have evolved had it not been for the generosity and active support of key researchers, clinicians, industry, and public institutions. It now needs the involvement of new researchers, clinicians, and entrepreneurs to continue its development.

## Acknowledgements

Veterans Administration Merit Review (I01RX-001821-01A1) and This work was supported by Computational Optometry (Atarfe, Spain, URL: www.optocom.es). We thank Jize Dong for writing the initial versions of the Java code for the OPI Daydream/Cardboard server. We also thank Zachary Heinzman for reviewing the manuscript.

